# Orthogonal Transposons for Iterative Genome Engineering of Mammalian Cells

**DOI:** 10.64898/2026.03.24.714049

**Authors:** Maggie Lee, Sowmya Rajendran, Divya Vavilala, Lynn Webster, Indira Kottayil, Ferenc Boldog, Mario Pereira, Maya Wright, Surya Karunakaran, Molly Hunter, Varsha Sitaraman, Claes Gustafsson, Jeremy Minshull

## Abstract

The contemporary shift toward multispecific antibodies, antibody-drug conjugates (ADCs), and bespoke glycoengineered therapeutics have exposed the limitations of standard genomic engineering tools. This paper presents a novel iterative engineering paradigm utilizing the Leap-In Transposase^®^ platform. By leveraging a suite of three mutually orthogonal transposase-transposon systems, we demonstrate the sequential modification of the Chinese Hamster Ovary (CHO) genome to achieve three distinct functional outcomes: (i) First, the creation of a glutamine synthetase (GS)-deficient host (CHO-K1-GS) via targeted knockdown, (ii) Second, the integration of multiple copies of a model therapeutic IgG1 for expression, and (iii) Third, the subsequent knockdown of the fucosylation pathway to modulate the glycan profile of the expressed IgG1. Genetic stability (copy number & sequence) of each integration event was confirmed using Targeted Locus Amplification (TLA) and Next-Generation Sequencing (NGS). Functional stability (expression levels, metabolic phenotype, and glycan phenotypes) was confirmed using standard cell culture and analytical techniques. Crucially, the truly orthogonal nature of the transposase-transposon pairs prevents cross-mobilization and ensures the structural and functional integrity of previously integrated cargo. This study establishes a “What You See Is What You Get” (WYSIWYG) methodology that provides a robust, scalable, and predictable framework for developing next-generation complex biopharmaceutical manufacturing cell lines.

## Introduction

Since the 1980s, Chinese Hamster Ovary (CHO) cells have been the predominant mammalian host cell line for the production of recombinant biotherapeutics. Their prominence is not incidental; CHO cells exhibit several critical industrial traits, including the ability to grow in high-density, chemically defined cultures and a post-translational modification (PTM) machinery that generates human-compatible N- and O-linked glycosylation profiles. Researchers continue to engineer better CHO hosts by targeting genes that improve growth characteristics, PTM machinery & metabolic profiles. While the need for improved expression hosts has evolved, available genome-engineering tools remain limited by problems such as the low efficiency of homology-directed repair in targeted nuclease-mediated genome editing, the need to screen genomic hotspots for ‘landing pad’ approaches extensively, and the lack of genomic stability observed with random integration approaches. In addition to this demand for better genome-editing tools, the complexity of biotherapeutic formats designed and evaluated for various diseases has also increased exponentially. We are no longer limited to small proteins or simple monoclonal antibodies (mAbs). The current biologics pipelines include an ever-evolving variety of bispecifics, trispecifics, Fc-fusion proteins, and complex vaccines. These molecules require precise control over gene dosage, chain ratios, and metabolic backgrounds — demands that have pushed traditional CHO engineering methods to their breaking point.

Transposons, or “jumping genes,” discovered by Nobel laureate Barbara McClintock (McClintock 1950), offer a compelling alternative path. Transposases catalyze a cut-and-paste mechanism that relocates discrete DNA segments flanked by Inverted Terminal Repeats (ITRs) into the host genome. Unlike targeted nucleases, engineered hyperactive transposase systems prioritize integration into open chromatin, *i*.*e*., transcriptionally active regions. Today, transposon-based technologies are increasingly dominating modern cell line development. Transposon-transposase pairs such as piggyBac (Ding *et al*. 2005), Sleeping Beauty (Ivics *et al*. 1997), and Leap-in Transposase® (Balasubramanian *et al*. 2018; Lee *et al*. 2019), are widely used in cell line development to accelerate the creation of stable, high-producing mammalian cells. These “cut-and-paste” DNA systems enable efficient, semi-targeted insertion of genes into active transcription sites, leading to higher titers and stable integration.

ATUM’s Leap-In Transposase^®^ platform has emerged as the gold standard in this space (Rajendran *et al*. 2021). It overcomes the limitations of random integration by ensuring that every integrated copy of the transgene is structurally intact and functional. By integrating multiple copies (typically 5–50) distributed across the genome, Leap-In Transposase rapidly creates highly productive, genetically stable cell populations. During the COVID-19 pandemic, several therapeutic antibodies were brought to clinical trials at unprecedented speed, in large part due to the Leap-In platform (Schmieder *et al*. 2022: Agostinetto *et al*. 2022). In addition to the transposase/transposon itself, the Leap-In platform also includes a portfolio of optimized vector elements (Rajendran *et al*. 2025; Wang et al. 2024; Wang *et al*. 2022), bespoke sequence optimization algorithms (Theodorou *et al*. 2026), and the CHO-K1-based host cell line miCHO.

However, using a single transposase system is limiting when iterative, sequential cell engineering steps are required. In a conventional system, expressing the same transposase for a second modification carries the risk of cross-mobilization, in which the transposase identifies and excises previously integrated transposons. To build a robust toolbox for complex cell engineering needs, we have developed five orthogonal transposon-transposase pairs derived from five different organisms. In nature, each transposase has co-evolved with a specific ITR sequence, enabling these orthogonal pairs to sequentially engineer host cell characteristics, such as improving sialylation, reducing fucosylation, or generating hosts with convenient selection markers. This study uses three of the five orthogonal transposases to demonstrate the power of orthogonal transposase-transposon pairs to sequentially engineer a wild-type CHO host for metabolic selection, followed by high-titer expression of an antibody biotherapeutic in the engineered host, and subsequent modulation of the antibody product quality (Fig 1a).

**Figure 1.**
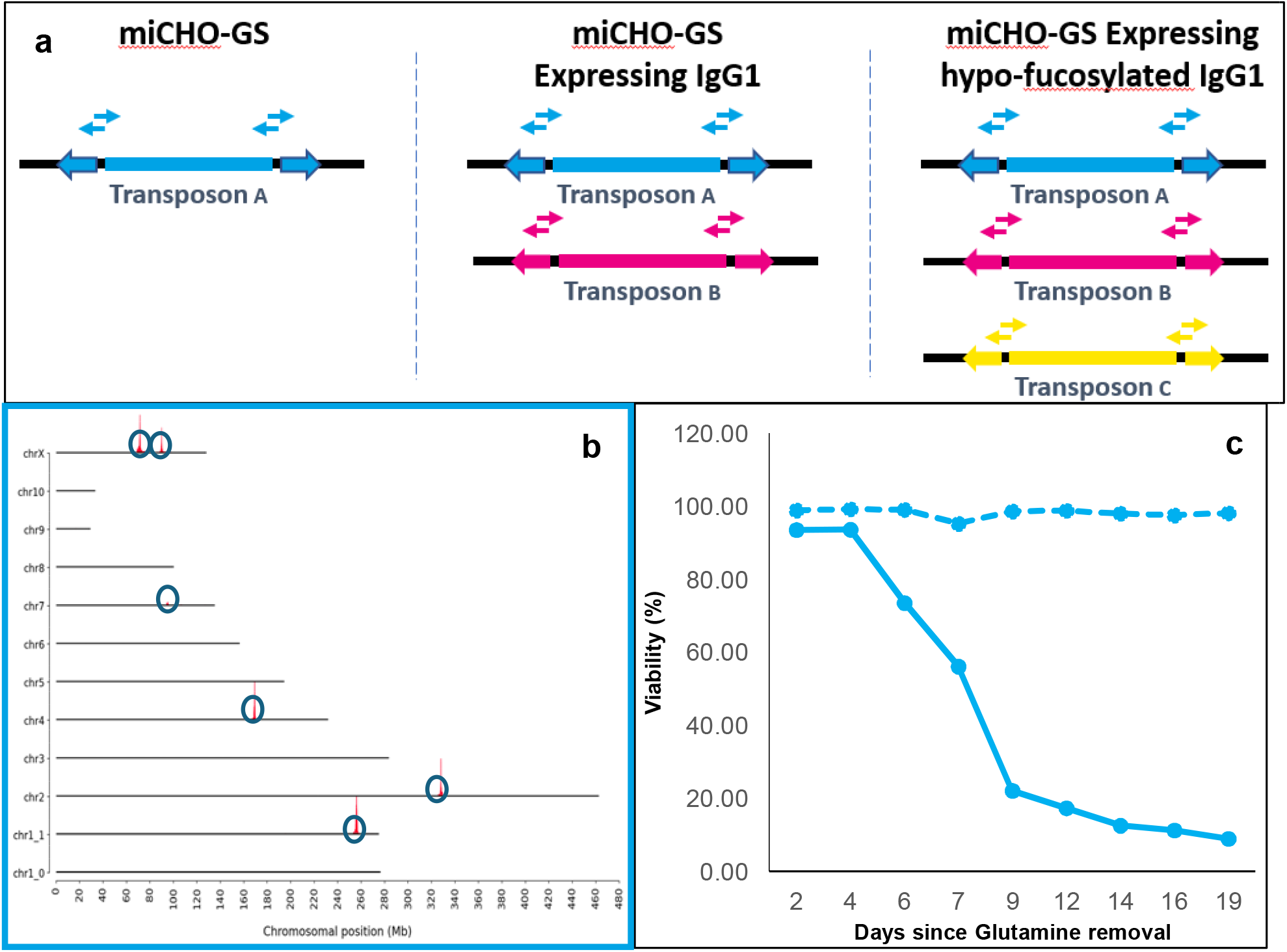
a) Schematic showing the experimental setup & TLA analysis strategy: Three orthogonal transposase-transposon pairs were used to engineer a CHO-K1 host cell line sequentially. Two transposon-specifi primer pairs were designed to confirm the number and stability of integrated copies by TLA analysis. b) Each horizontal line represents a CHO chromosome, as denoted on the left. Integrants are mapped by TLA to the corresponding chromosomal location as peaks, circled in blue for transposon A integration events. Six independent transposon A integration sites are identified in the resulting clonal miCHO-GS cell line. c) Viability of miCHO host with transposon A (for Glutamine Synthetase knockdown). The transposon A-transfected miCHO (solid blue line) does not survive in glutamine-free media and requires glutamine addition (dotted blue line), demonstrating successful knockdown of the endogenous GS.

## Materials and Methods

### Transposase and vector designs

The three engineered Leap-In transposases A, B, and C used in this study are derived from three different host organisms and share only minimal sequence homology. All three transposase enzymes were engineered and optimized using ATUM’s machine-learning platform (Liao *et al*. 2007) to achieve transposition efficiency significantly higher than that found in nature. Each transposon vector (A, B, and C) was constructed with mutually orthogonal ITR sequences, ensuring that the cognate B transposase would not recognize or excise the integrated elements from transposase A. The transposon vectors utilize ATUM’s proprietary modular architecture, including optimized promoters, untranslated regions (UTRs), signal peptides, and mRNA transport sequences. Codon optimization was performed using ATUM’s proprietary algorithms (Welch *et al*. 2011) to maximize translational efficiency while avoiding mRNA secondary structures and other sequence features that may impede expression.

### Cell line, culture conditions, transfections, and analytics

The parental CHO-K1 cell line was used as the host for the first set of transfections to knock down the endogenous glutamine synthetase gene. A single-cell clone from this engineered host pool was used as the host for the second transfection to express the antibody sequence. A single cell clone identified from the antibody-expressing pool was used as the host for the third transfection to knock down fucosylation. Cells were maintained in chemically defined, animal-component-free media in high-density suspension cultures. Standard incubation conditions were 37°C, 5% CO_2_, and 120 rpm agitation. Co-transfection of the transposon vector encoding the gene(s) of interest and mRNA encoding the Leap-In Transposase was performed using standard procedures. Cell growth and viability were monitored periodically by automated cell counters. IgG1 titers were determined via Protein A HPLC. The glycan profile of the secreted antibodies was analyzed using hydrophilic interaction chromatography (HILIC). Genetic stability was assessed by maintaining the final clones in continuous culture for 60 generations and re-evaluating integrated transgene copy numbers, clone productivities, and glycan profiles.

### Genetic characterization by TLA

To confirm the precision and stability of the integrations, we utilized Targeted Locus Amplification (TLA) in partnership with Solvias (Utrecht). TLA is a proximity ligation-based method that enables the complete sequencing of a transgene and its surrounding genomic environment without prior knowledge of the integration site (de Vree *et al*. 2014; Cain-Hom *et al*. 2017). Two sets of transposon-specific primers were designed for each of the three systems to: (i) map the precise chromosomal coordinates of every integration, (ii) count the number of copies integrated, (iii) verify the sequence and structural integrity of the cassettes (absence of concatemers or truncations), and (iv) monitor for any cross-mobilization or rearrangements during sequential steps.

## Results

### Engineering of the miCHO-GS metabolic host

The primary objective of the first transposon transfection was to engineer a wild-type CHO-K1 cell line into a glutamine synthetase-deficient cell line without using traditional nuclease-dependent genome-editing methods. Leap-In transposon A, which encodes genetic elements to knock down the endogenous GS, was co-transfected using standard procedures with an mRNA encoding transposase A. Single cell clones from the stable pool were selected based on growth, productivity, and transfectability, and the lead clone was identified as the subsequent miCHO-GS host (engineered host where endogenous GS was knocked down). Functional characterization of the miCHO-GS host clearly showed that it cannot grow in the absence of glutamine in the culture medium, demonstrating successful knockdown of the glutamine synthetase gene (Fig. 1c). The TLA analysis of the clonal miCHO-GS host identified 6 distinct integration loci for transposon A (Fig. 1b). These 6 integration events were found to be stable for ~75 generations. The resulting miCHO-GS host generated in step 1 was subsequently transfected with transposon and transposase B, a transposon which encodes both an IgG1 antibody and the glutamine synthase gene. The stable integration of transposon B rescued the GS-deficient phenotype by providing a recombinant GS gene alongside the IgG1 heavy and light chains. Single-cell clones were isolated from the stable miCHO-GS pool expressing the antibody. Subsequent TLA analysis of the lead clone revealed transposon B integrations at 33 genomic loci (Fig. 2a). Phenotypic validation of the resulting cell line was unequivocal: the miCHO-GS host failed to survive in glutamine-free media, demonstrating a successful functional knockdown of the endogenous GS enzyme, whereas the transposon A+B cell line showed full rescue at day 6 (Fig. 2b). The resulting cell pools showed rapid recovery and high productivity, with antibody titers exceeding several grams per liter in standard laboratory fed-batch processes (Fig. 2c). A derivative transposon A + transposon B clonal cell line was expanded for an additional ~75 generations (a total of ~150 generations for the transposon A integrations). Despite the high number of integrations, the structural integrity of each transposon A and transposon B copy was perfectly maintained nucleotide by nucleotide, as confirmed by next-generation sequencing (data not shown). Importantly, TLA analysis confirmed that the 6 original integration sites from transposon A remained undisturbed, demonstrating the strict orthogonality of the two transposase systems.

**Figure 2.**
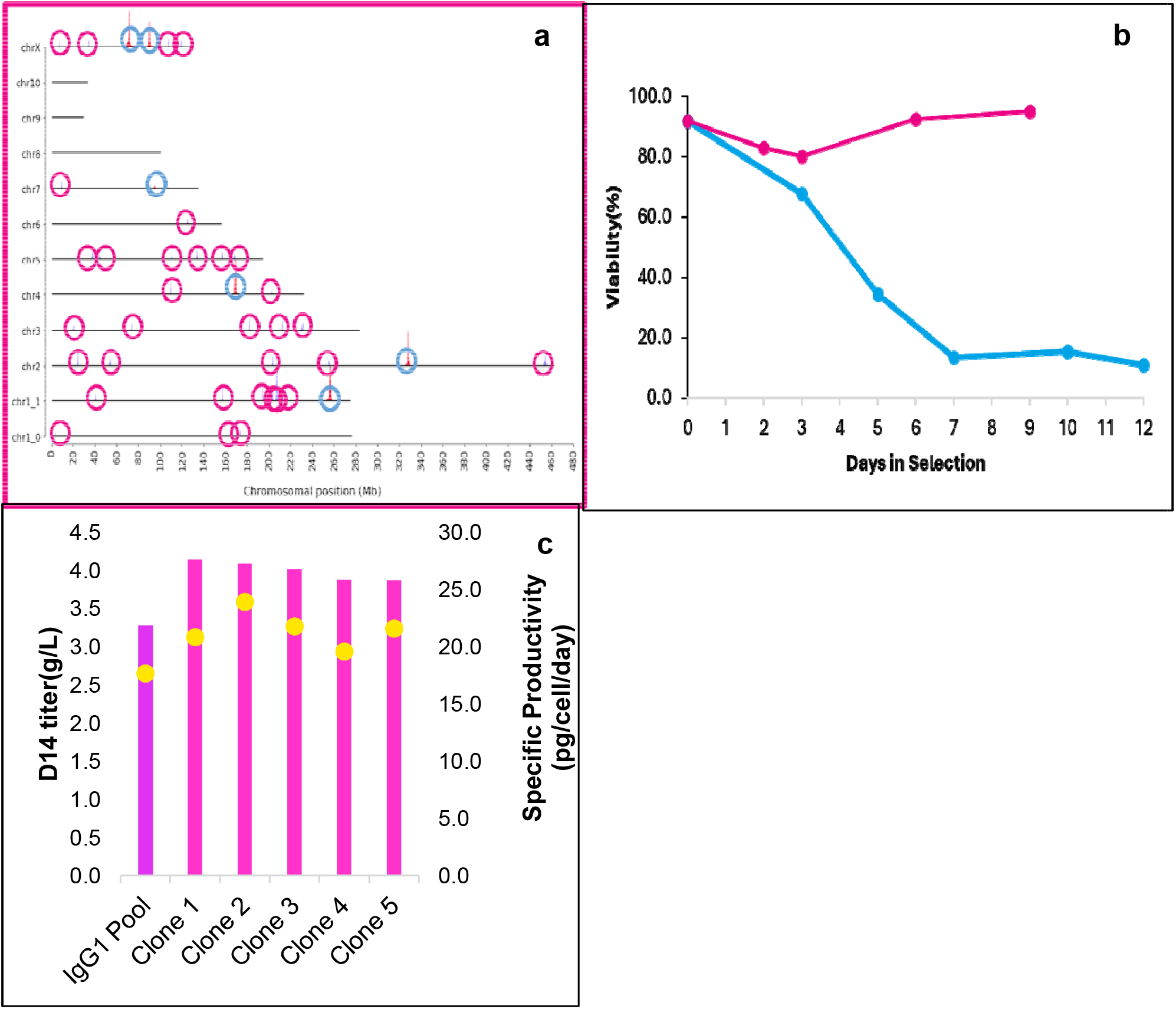
a) Each horizontal line represents a CHO chromosome, as denoted on the left. Integrants are mapped by TLA to the corresponding chromosome as peaks, circled in blue for transposon A integration events, and circled in pink for transposase B integration events. Six transposon A integration site and 33 independent transposon B integration sites are identified in the resulting clonal miCHO-GS-mAb cell line. b) Viability of miCHO with transposon A, with or without transposon B transfection. The transposon A-transfected miCHO (blue) does not survive in glutamine-free media, demonstrating successful knockdown of the endogenous GS. Transfection of miCHO-GS with transposon B encoding an antibody and the GS gene (pink) shows full rescue by day 6. c) Antibody titer of the miCHO-GS pool after transfection of B results in ~3.3g/L, and the top clones 1-5 derived from the pool produce between 3.1 and 4.3g/L. Specific productivity of the top clones is between 18 and 24 pg/cell/day (yellow dots)

### Glyco-engineering for enhanced antibody-dependent cell-mediated cytotoxicity (ADCC)

To further demonstrate iterative cell line engineering, the miCHO-GS host (engineered by stably integrating transposon A to knock down glutamine synthetase expression) expressing the antibody sequences (transposon B integrations) was transfected with transposon C. This transposon C encodes genetic elements that target and knock down FUT8 fucosyltransferase in the fucosylation pathway. The TLA analysis of Transposon C expressing miCHO-GS-mAb clone identified 9 integration sites for Transposon C (Fig. 3a). As before, the 6 integration sites from transposon A and the 33 from transposon B remained perfectly intact. The resulting triple-engineered cell line expressed IgG1 at a similarly high titers to the parental miCHO-GS-mAb clone (transposon A+B cell line) (Fig. 3b), but with an altered glycan profile consistent with knockdown of the fucosylation pathway (Fig. 3c).

**Figure 3.**
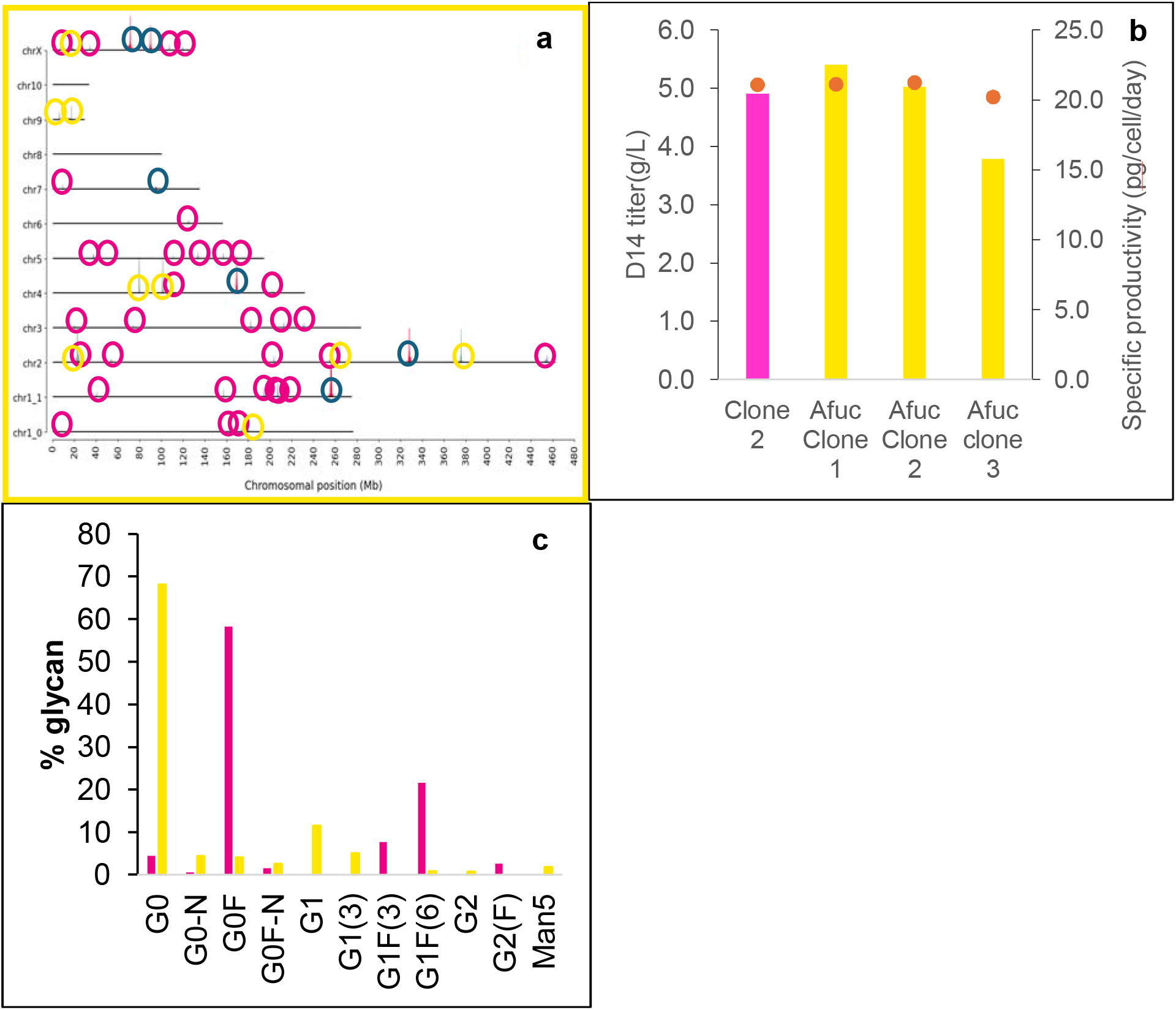
a) Each horizontal line represents a CHO chromosome, as denoted on the left. Integrants are mapped by TLA to the corresponding chromosome as peaks, circled in blue for transposon A integration events, circled in pink for transposase B integration events, and circled in yellow for transposon C integration events. Six transposon A integration sites, 33 transposon B integration sites, and 9 transposon C integration sites are identified in a derivative clonal miCHO-GS-mAb-afuc cell line. b) IgG1 productivity from miCHO-GS-mAb (transposon A+B pool) ~5g/L (pink bar). Afucosylated IgG1 productivity of the top 3 clones (yellow bars) after additional transfection with transposon C that knocks down fucosylation *fut8* ranges from 4.7 to 5.3 g/L (~20 pg/cell/day, denoted as red dot). No reduction in recombinant antibody titer was observed in the miCHO-GS-mAb-afuc (transposon A+B+C) cell line compared to the parental miCHO-GS-mAb (transposon A+B) cell line. c) Pink bars denote the fraction of each glycan species observed in the clone transfected with transposons A+B. Yellow bars represent the fraction of each N-linked glycan species observed in the clone transfected with transposons A+B+C. N-linked glycan structures are designated as follows: G0, G1, G2, indicating 0, 1, or 2 terminal galactose residues, respectively. F indicates the presence of core fucose. N represents the presence of bisecting N-acetylglucosamine (GlcNAc). Man5 denotes an oligomannose structure consisting of 5 mannose residues on the 2 N-acetylglucosamine (GlcNAc) base.

Analysis of the IgG1 glycan species showed a substantial shift in glycosylation profile. Total fucose levels were reduced to under10%, with a corresponding increase in non-fucosylated G0, G1, and G2 species (Fig. 3c).

The triple-transfected miCHO-GS-mAb-afuc cell line was expanded for another ~90 generations, for a total of 240 generations (75 generations after the first transfection, 75 generations after the second transfection, and 90 generations after the third transfection). Stability studies showed no significant loss in specific productivity or volumetric productivity with the addition of transposon C. The hypo-fucosylation phenotype (under 10% fucosylation) remained stable throughout the stability study. Furthermore, TLA analysis at the end of 90 generations confirmed that all 48 integration sites (6 of transposon A, 33 of transposon B, and 9 of transposon C) remained at their original chromosomal positions with no sequence alterations, as shown in Fig. 3a.

## Discussion

### The “WYSIWYG” Paradigm in Biotechnology

Transposon-mediated genome engineering validates a “What You See Is What You Get” (WYSIWYG) paradigm for cell line engineering. In traditional cell line engineering, the genetic elements to be inserted (the vector and the genetic cargo) are often altered by the cell’s repair machinery, leading to insertions, deletions, truncations, concatemerizations, junction mutations, and other unpredictable outcomes. In contrast, the Leap-In platform ensures that the genetic information is transferred 1-to-1 from the vector to the host genome. What is designed on the desktop is consistent with the genetic information encoded in the cell, both at the initial pool stage and later in the clonal cell line. The Leap-In platform provides genome-editing tools for the complex, iterative engineering of mammalian hosts. The genomic and functional stability afforded by the Leap-In platform is illustrated here through three successive transfections with three orthogonal Leap-In transposon-transposase pairs, and by assessing stability for at least 60 generations to demonstrate the genetic stability of each generated cell line. All transposon integration sites from the three serial transfections are stable, and any phenotypic drift is below the detection limit. This stability allows, for example, IND-enabling studies using pool-derived material, saving significant time in the drug development lifecycle (Jiang et al. 2025; Pan et al. 2025).

### Orthogonality as an enabling tool for exploring genetic complexity

The use of orthogonal transposase-transposon pairs represents a new paradigm in iterative CHO engineering. Successive engineering steps are often required to express complex biologics or to modulate PTMs. A toolbox with orthogonal transposases can be used to build multichain molecules, e.g., by integrating two chains using a single transposon, then introducing a third or fourth chain using an orthogonal system to determine the optimal chain ratio for the expression of complex multichain molecules. An alternative application, as shown here, is the independent tuning of metabolic pathways (GS) and glycosylation pathways relative to the therapeutic protein. Orthogonal transposons can accordingly be used to retroactively engineer an existing clonal cell line to introduce properties in the protein therapeutic, enabling new clinical indications. We also anticipate the development of an ever-expanding library of bespoke hosts with different protease, glycosylation, and other metabolic and regulatory patterns. Orthogonal transposases can be used serially, as illustrated here, or in multiplex format to explore multiple alternative genome-engineering strategies simultaneously.

### Regulatory and CMC Implications

The long-term genetic stability demonstrated over three serial transpositions is a critical requirement for regulatory approval (IND/BLA). The ability of TLA analysis to rapidly map the exact genomic location of every transgene across multiple iterations provides critical product quality monitoring and molecular characterization of the drug-producing cell line, and de-risks the CMC process. To date (February 2026), there have been 50 INDs for molecules produced via Leap-In technology across the US, EU, Australia, and China. This clinical track record proves that the platform is not just a research tool but a validated manufacturing engine.

Our study demonstrates a versatile toolbox where a combination of hyperactive, orthogonal transposases and modular synthetic biology approaches can be used to tackle the most challenging problems in CHO cell engineering. We have shown that it is possible to sequentially and stably alter multiple metabolic and product-quality pathways while maintaining high protein expression levels. The Leap-In platform’s ability to generate high-titer, stable, and well-characterized cell lines rapidly addresses the industry’s most pressing need: speed-to-clinic. As we move into an era of increasingly complex biologics, the robustness and predictability of orthogonal transposon systems will be the foundation for the next generation of life-saving therapies.

## Acknowledgments

The authors would like to thank the team at Solvias NL for their invaluable support in performing the TLA analysis. We also thank the ATUM bioprocessing team for their work in fed-batch characterization and glycan analysis. ATUM fully funded the work.

## References

McClintock, B. (1950). The origin and behavior of mutable loci in maize. Proc Natl Acad Sci USA. 36(6):344–55

Ding S, et al. (2005). Efficient transposition of the piggyBac (PB) transposon in mammalian cells and mice. Cell. 122(3):473–83

Ivics Z, et al. (1997). Molecular reconstruction of Sleeping Beauty, a Tc1-like transposon from fish, and its transposition in human cells. Cell. 91(4):501–10

Balasubramanian et al. (2018). Generation of high-expressing Chinese hamster ovary cell pools using the Leap-In transposon system. Biotechnol J.13(10):e1700748

Lee M, et al. (2019). Leap-In transposases and transposons accelerate cell line development. Genet Eng Biotechn N. 39(9):59–61.

Rajendran S, et al. (2021). Accelerating and de-risking CMC development with transposon-derived manufacturing cell lines. Biotechnol Bioeng. 119(4):1041–52.

Schmieder V, et al. (2022). Towards maximum acceleration of monoclonal antibody development: Leveraging transposase-mediated cell line generation to enable GMP manufacturing within 3 months using a stable pool. J Biotechnol. 10(349):53–6

Agostinetto et al. (2022). Rapid cGMP manufacturing COVID-19 monoclonal antibody using stable CHO cell pools. Biotechnol Bioeng. 119(2):663–6

Rajendran S, et al. (2025). Cell line development for bispecific antibodies: better predictability through transposases. BioRxiv doi.org/10.1101/2025.08.12.669435

Wang Y, et al. (2024). Mitigating the aggregation challenge in immunocytokine production: Strategies during cell line development and purification optimization. Biochem Eng J. doi.org/10.1016/j.bej.2024.109215

Wang Y, et al. (2022). An innovative platform to improve asymmetric bispecific antibody assembly, purity, and expression level in stable pool and cell line development. Biochem Eng J. doi.org/10.1016/j.bej.2022.108683

Theodorou E, et al. (2026). Machine learning method for optimizing coding sequences in mammalian cells. bioRxiv doi.org/10.64898/2026.01.26.701778

Liao J, et al. (2007). Engineering proteinase K using machine learning and synthetic genes. BMC Biotechnol. 7:16

Welch M, et al. (2011). Designing genes for successful protein expression. Methods Enzymol. 498:43–66

De Vree PJP, et al. (2014). Targeted sequencing by proximity ligation for comprehensive variant detection and local haplotyping. Nat Biotechnol. 32(10):1019–25

Cain-Hom C, et al. (2017). Efficient mapping of transgene integration sites and local structural changes in Cre transgenic mice using targeted locus amplification. Nucleic Acids Res. 45(8):e62

Jiang Y, et al. (2025). Accelerating IND-enabling toxicology studies using protein products from stable pools or pools of clones in Chinese hamster ovary cells. Biotechnol Prog. 41(5):e70040

Pan J, et al. (2025). Utilizing non-clonal CHO cell derived materials for preclinical studies of complex molecules. BMC Biotechnol. 25(1):33

